# Vimentin supports cell polarization by enhancing centrosome function and microtubule acetylation

**DOI:** 10.1101/2023.02.17.528977

**Authors:** Renita Saldanha, Minh Tri Ho Thanh, Nikhila Krishnan, Heidi Hehnly, Alison Patteson

## Abstract

Cell polarity is important for controlling cell shape, motility, and cell division processes. Vimentin intermediate filaments are necessary for proper polarization of migrating fibroblasts and assembly of vimentin and microtubule networks is dynamically coordinated, but the precise details of how vimentin mediates cell polarity remain unclear. Here, we characterize the effects of vimentin on the structure and function of the centrosome and the stability of microtubule filaments in wild-type and vimentin-null mouse embryonic fibroblasts (mEFs). We find that vimentin mediates the structure of the pericentrosomal material, promotes centrosome-mediated microtubule regrowth, and increases the level of stable acetylated microtubules in the cell. Loss of vimentin also impairs centrosome repositioning during cell polarization and migration processes that occur during wound closure. Our results suggest that vimentin modulates centrosome structure and function as well as microtubule network stability, which has important implications for how cells establish proper cell polarization and persistent migration.

## Introduction

Animal cells must be dynamic to move and change shape while also stable and rigid to generate sustained polarized motion. Central to these features is the cell’s cytoskeleton. The animal cell’s cytoskeleton is comprised of three interconnected filamentous networks: F-actin, microtubules, and intermediate filaments (*1*). These networks coordinate different functional aspects of the cell. Filamentous actin is the cell’s main force-generating machinery, producing protrusive forces at the cell membrane and working with myosin motors to generate cellular contractile forces (*2, 3*). Microtubules are rigid polarized polymers that span the length of cell and direct cargo transport and organization within the cell (*4-6*), and intermediate filaments form a passive filamentous network that supports and stabilizes the cell (*7-10*). Many cellular functions rely on coordinated crosstalk amongst these cytoskeletal networks, though there is much more known about how they work individually than together (*11, 12*).

Several emerging studies point to a mutually reinforcing connection between vimentin intermediate filaments (IF) and the microtubule network (*9, 13-15*). Vimentin is an IF protein expressed in mesenchymal cells and highly invasive cancer cells (*16-18*). Disruption of microtubules collapse the vimentin network (*13*) and likewise disrupting vimentin alters the microtubule network (*15*). Vimentin has been shown to enhance the persistence of the microtubule network by serving as a stable long-standing template for new microtubule growth (*9*), and recent work has shown in reconstituted *in vitro* systems that vimentin stabilizes microtubules against depolymerization through direct interaction (*14*). Together, these works highlight a subtle interplay between microtubules and vimentin that is important for polarized cell migration.

The cell’s centrosome acts as the microtubule-organizing center of the cell and serves as the main microtubule-nucleating organelle (*19*). The centrosome is essential for whole-cell polarization and is typically positioned near the cell nucleus between the nucleus and the leading edge of the cell (*20*). Vimentin IFs are also localized around the cell nucleus, forming a dense physical mesh in the perinuclear region of the cell (*7, 10, 16, 17*). Experiments with cells on patterned adhesive substrates show loss of vimentin increases variability in centrosome positioning (*21*). Cells lacking vimentin also have impaired polarization (*22-26*), which suggest that vimentin might interact with both microtubules and their organizing center, the centrosome.

Based on these studies, we hypothesized a functional link between vimentin and the cell’s centrosome in establishing cell polarization and directed migration. Here we report novel experimental data that addresses the role of vimentin intermediate filaments in centrosome structure and microtubule-nucleating function. Our results suggest new mechanisms by which vimentin mediates cell polarization and cross-talks with the cell’s microtubule architecture.

## Materials and Methods

### Cell Culture

Wild-type (vim^+/+^), vimentin-null (vim ^-/-^) mouse embryonic fibroblasts (mEFs) were kindly provided by J. Ericsson (Abo Akademi University, Turku, Finland). Cells were maintained in Dulbecco’s Modified Eagle’s Medium (DMEM) including HEPES and sodium pyruvate supplemented with 10% fetal bovine serum (FBS), 1% penicillin streptomycin, and non-essential amino acids. Cell cultures were maintained at 37°C with 5% CO_2_.

### Immunofluorescence

Cells were fixed for immunofluorescence using methanol for 10 minutes at -20°C. Cell membranes were permeabilized with 0.05% Triton-X in PBS for 15 minutes at room temperature and blocked with 1% bovine serum albumin (BSA) for 1 hour at room temperature. For vimentin visualization, cells were incubated with primary rabbit anti-vimentin antibody (Abcam) diluted 1:200 in 1% BSA in PBS for 1.5 hour at room temperature; the secondary antibody anti-rabbit Alexa Fluor 555 (Invitrogen) was used at a dilution of 1:1000 in 1% BSA in PBS for 1 hour at room temperature. For visualizing microtubules, we used primary alpha-tubulin rat antibody (Bio-rad) diluted 1:200 in 1% BSA in PBS for 1.5 hours at room temperature and secondary antibody anti-rat Alexa Fluor 647 (Invitrogen) at a dilution of 1:1000 in 1% BSA in PBS for 1 hour at room temperature. For visualizing acetylated microtubules, cells were incubated with primary acetylated tubulin mouse antibody (Sigma Aldrich) diluted 1:200 in 1% BSA in PBS and incubated for 1.5 hour at room temperature; secondary antibody anti-mouse Alexa Fluor 568 (Invitrogen) was used at a dilution of 1:1000 in 1% BSA in PBS for 1 hour at room temperature. Cells were stained using Hoechst 33342 (Molecular Probes) for 1 hour at room temperature. For visualizing centrosome, cells were incubated with primary Cdk5rap2 rabbit antibody (Bethyl Laboratories), gamma-tubulin rabbit antibody (Sigma Aldrich), Pericentrin rabbit anibody (Abcam), Cenexin rabbit antibody (Protein tech) and centrin mouse antibody (EMD Millipore) diluted 1:1000 for centrin and 1:200 for other centrosomal proteins in 1% BSA in PBS for 1.5 hours at room temperature; secondary antibody anti-mouse Alexa Fluor 488 (Invitrogen) for centrin and anti-rabbit Alexa Fluor 488 (Invitrogen) for other centrosomeal proteins was used at a dilution of 1:1000 in 1% BSA in PBS for 1 hour at room temperature.

Cells were mounted using Prolong diamond antifade mountant (Life Technologies) for epi-fluorescence and confocal Imaging.

### Expansion microscopy

Cells were plated on 22x22 mm coverslips until they reached 90% confluence and fixed with ice cold methanol at –20°C for 10 mins followed by the immunofluorescence procedure mentioned above. The cells were stained with antibodies against vimentin and centrin. Expansion microscopy was performed using techniques similar to previously published protocols (*27, 28*). Briefly, the fixed cells were then incubated at 40° C overnight in incubation solution (30% acrylamide in 1X PBS). Next, 1 ml of freshly prepared gelation solution (20% acrylamide, 7% sodium acrylate, 0.04% bis acrylamide, 0.5% APS, and 0.5% TEMED in 1× PBS) was added per coverslip and allowed to solidify on ice for 20 mins, followed by incubation for 20 mins at room temperature and additional 1.5 hours at 30°C. The solidified gels were then sectioned into 4 mm gel punches using a disposable biopsy punch.

The gel punches were then digested with digestion buffer overnight (0.5% Triton-X, 0.03% EDTA, 1M Tris–HCl, pH 8, 11.7% sodium chloride and 1U/ml of proteinase K). Finally, the punches were subjected to a second round of immunofluorescence procedures with antibody concentration increased to twice of the initial concentration and allowed to expand in water overnight at 4°C. The expanded gel punches were then mounted on a 35-mm Mattek dish and imaged using Leica SP8 confocal microscope.

### Western Blot

Cell lysates were obtained by suspending the cells in lysis buffer (HSEG buffer pH 7.4 - 40 mM NaCl (Fischer Scientific), 5mM EDTA (Fischer Scientific), 4% Glycerol (Fischer Scientific), 20mM NaF (Fischer Scientific), 1% TritonX-100 (Fischer Scientific), 1X protease inhibitor; 0.1mM PMSF). Bio-Rad Protein Assay Kit II (Bio-Rad laboratories) was used to measure the acetylated tubulin concentration from the post-nuclear supernatant collected from the lysates. Standard Western blot procedures were performed. The nitrocellulose membranes were probed with primary acetylated tubulin mouse antibody (Sigma-Aldrich) and primary alpha-tubulin mouse (Sigma-Aldrich) antibody diluted in TBS-Tween20 (ThermoFischer) and incubated overnight at 4^0^C. The membranes were probed using mouse horseradish peroxidase conjugated secondary antibody (Invitrogen) for an hour at room temperature. The protein levels were visualized using Clarity ™ western ECL (Bio-rad Laboratories) substrate and imaged using Bio-Rad ChemiDoc ™ imager.

### Microtubule re-nucleation experiments and microtubule analysis

Cells were first treated with 1μM nocodazole (Sigma-Aldrich) for 30 minutes at room temperature to disrupt microtubule filaments. To allow for microtubule regrowth, nocodazole was washed out and cells were placed back in cell media. Cells were fixed at 0, 1, 2, and 5 minute intervals after washout and stained for microtubules and vimentin. The microtubule re-growth was analyzed by tracing the radial region of microtubules stemming from the cell centrosome. Tracing was done manually in FIJI-ImageJ software. A minimum of 75 cells were analyzed over 3+ independent experiments per condition. For analyzing microtubules after nocodazole treatment, we used the Source Steger’s algorithm in curve trace (*29*) in FIJI-ImageJ and computed the total microtubule contour length per cell; a minimum of 90 cells were analyzed over 3+ independent experiments per condition.

### Scratch Wound-Healing Assay

A monolayer of cells was created by culturing 5x10^5^ cells in a 35-mm glass bottom petri dish overnight. Wounds were generated using a 10 μl pipette tip to scratch the monolayer. Following the scratch wound, cells were fixed at different time points (1, 2 and 4 hrs) post scratching. The monolayers were then stained for centrosome (cdk5rap2), microtubules (alpha-tubulin), and DNA (Hoechst 33342) and imaged using a Nikon epi-fluorescence microscope. Cells within 20μm of the scratch were analyzed for the centrosome position. Centrosome positioning with respective to the wound was analyzed using FIJI-ImageJ. A minimum of 70 cells were analyzed over 2+ independent experiments per condition.

### Imaging

#### Epi-fluorescence imaging

Epi-fluorescence imaging was performed using a Nikon Eclipse Ti (Nikon Instruments) inverted microscope equipped with an Andor Technologies iXon em+ EMCCD camera (Andor Technologies). Cells were imaged using a Plan Fluor (NA of 1.49) 100X oil immersion objective.

### Confocal imaging

Expansion microscopy images of centrosome and vimentin were obtained by using SP8 laser scanning confocal microscope with lightning equipped with HC PL APO 40×/1.10 W CORR CS2 0.65 water objective. Airy-Scan confocal imaging was used to image microtubules and acetylated microtubules. Images were obtained using Zeiss Airy-Scan LSM 980 confocal microscope equipped with Plan Apochromat 63X Oil immersion objective with 1.4 NA, 8Y multiplex mode and GaAsP detector. Images obtained were processed using Airy-scan processing technique from Zeiss Zen 3.2 software. For the Scratch assay, zoomed out images at 10x magnification were obtained using spinning disk confocal microscope (Yokogawa CSU-W1) on ab inverted Nikon Ti-E microscope with Perfect focus imaged onto a Andor Zyla CMOS camera.

### Statistics

Data are presented as a mean value ± standard error of mean (SEM). The unpaired Student’s t-test with 95% confidence level was used to determine statistical differences between distributions. Denotations: *, *p* ≤ 0.05; **, *p* ≤ 0.01; ***, *p* ≤ 0.001; ****, p ≤ 0.0001; n.s., *p* > 0.05. N represents number of independent experiments. n represents the number of cells. All the graphs and statistical analysis was done using GraphPad PRISM version 9 (GraphPad Software, Inc.).

## Results

### Loss of vimentin perturbs the pericentriolar matrix

We used laser scanning confocal and fluorescence microscopy to examine the centrosome in and vim ^-/-^ mEFs (Fig. 1a). To label the centrosome, cells were fixed and stained for the centrosomal protein Cep215 (cdk5rap2), a major pericentriolar material (PCM) protein associated with the organization of microtubules from the centrosome (*30, 31*). Figure 1a shows immunofluorescence images of microtubules, Cep215, and vimentin taken by a confocal laser scanning microscope. In wild-type cells, as expected, there is a strong accumulation of tubulin at the centrosome with long microtubule filaments extending out radially. Interestingly, in vim ^-/-^ cells, there is less tubulin localized around the centrosome (Fig 1b). Further, the centrosome itself (marked by Cep215) is noticeably more condensed and smaller in the vim ^-/-^ mEFs compared to vim^+/+^ mEFs. In addition, we observe an accumulation of vimentin that colocalizes with the centrosome in wild-type cells (Fig 1a)

**Figure 1:**
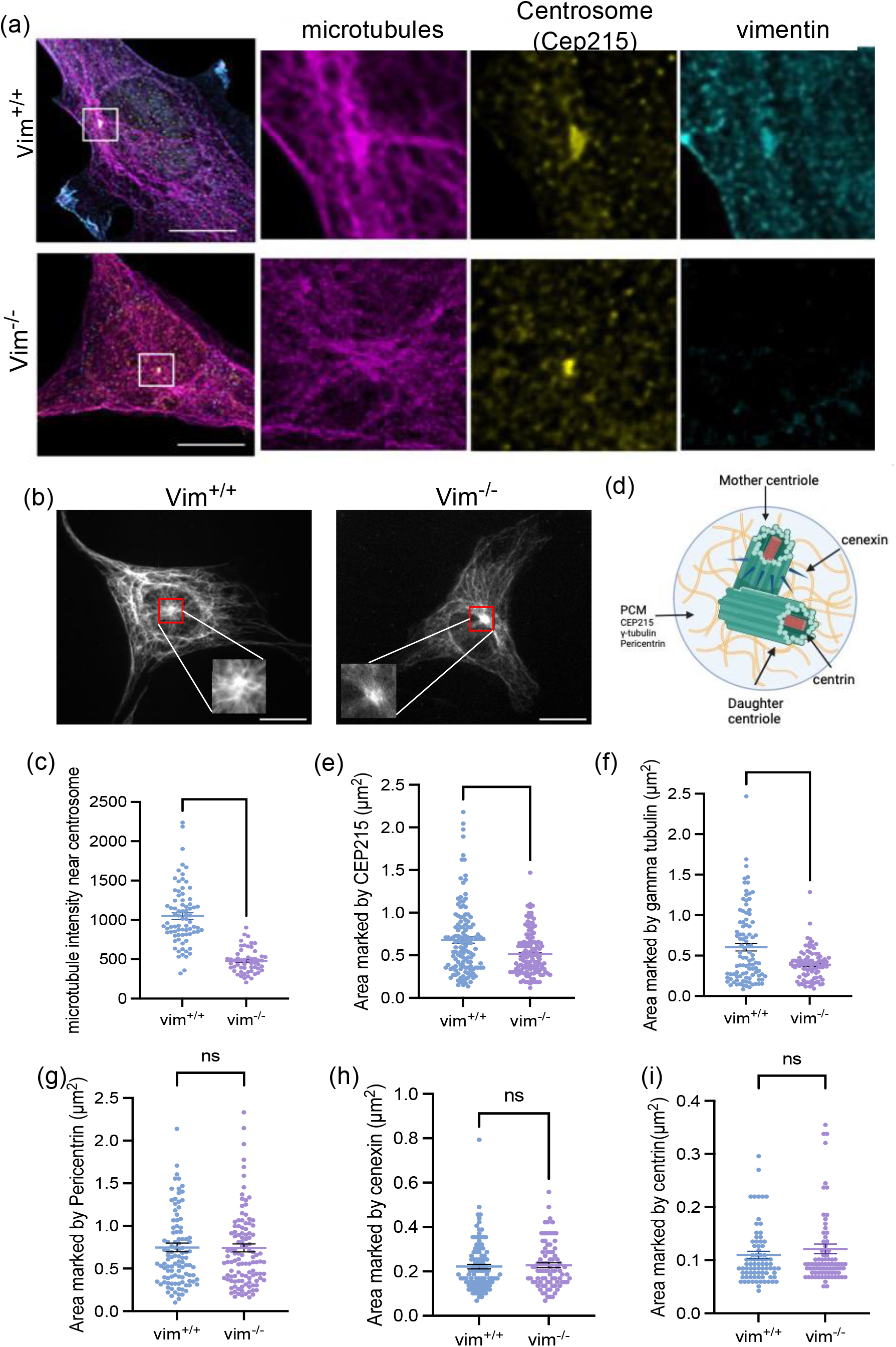
(**a**) Laser Scanning confocal images of microtubules, centrosome (CEP215) and vimentin in vim^+/+^ and vim^-/-^ mEF’s. (**b**) Epi-fluorescence microscopy images of microtubules originating from the centrosomes in vim^+/+^ and vim^-/-^ mEF’s . The insets show zoomed in images at the centrosome. **(c)** quantification of microtubule intensity near the centrsome (marked as a red box in vim^-/-^ cells) **(d)** Schematic of centrosome structure showing proteins associated with PCM and centrioles. **(e-i)** The projected area of the centrosome marked by CEP215 and gamma tubulin, pericentrin, cenexin and centrin. Denotation: ***, *p* ≤ 0.001; **, p ≤ 0.01) n=3, N>90 cells analyzed per condition. Scale bar is 20 μm.

Based on these observations, we next sought to quantify centrosome structure in vim^+/+^ and vim^-/-^ mEFs (Fig. 1a). The centrosome is composed of two centrioles (a mother and daughter centriole) embedded in a dense matrix of PCM proteins (Fig. 1d, schematic). To examine the PCM and centriole structure, we fixed and labeled cells for multiple centrosomal proteins localized in either the PCM (Cep215, γ-tubulin, pericentrin) or centriole protein complexes (cenexin and centrin). We found that the loss of vimentin significantly decreased the mean projected area of the centrosome by 40% as marked by Cep215 (Fig. 1e, p ≤ 0.001) and by 40% as marked by gamma-tubulin (Fig. 1f, p ≤ 0.001), which indicates that loss of vimentin perturbs the peri-centriolar matrix. There was no significant difference in centrosomal area observed by staining for pericentrin (Fig. 1g), but the intensity for cells lacking vimentin is lower compared to wild-type cells (p<0.001, SI Fig. 1c), suggesting cells lacking vimentin have less pericentrin protein localized to the centrosome. There was no significant difference observed for the mean area and intensity of centrosome marked by centriole proteins, cenexin (Fig 1h) and centrin (Fig. 1i). From these observations, we conclude that loss of vimentin disrupts multiple PCM proteins localization but does not affect centriole associated proteins, cenexin and centrin.

Since our data indicated loss of vimentin alters centrosome structure, we further analyzed centrosome structure using expansion microscopy (Methods). The centrioles are approximately 200nm in diameter, just within the spatial limit attainable by light microscopy (200-500 nm).

Expansion microscopy overcomes this barrier by physically increasing a specimen size by embedding a specimen in a swellable polymer matrix. By swelling the samples, the physical distance between proteins in the cell increases, allowing smaller structures to be resolved by light microscopy.

Figure 2 shows representative expansion microscopy images of wild-type and vimentin-null cells labeled with antibodies against vimentin and centrin. In wild-type cells, two distinct centrioles are observable in the expansion microscopy images. An accumulation of vimentin is also observed in the proximity of the centrioles. In contrast, two distinct centrioles are not observable in most of the vimentin-null cells, suggesting the centrioles in vimentin-null cells are much closer together and not spatially resolved even after expansion microscopy. These results further indicate a role of vimentin in maintaining centrosome structure that could be related to supporting the PCM (Fig. 2) and maintaining the spatial distance between centrioles.

**Figure 2:**
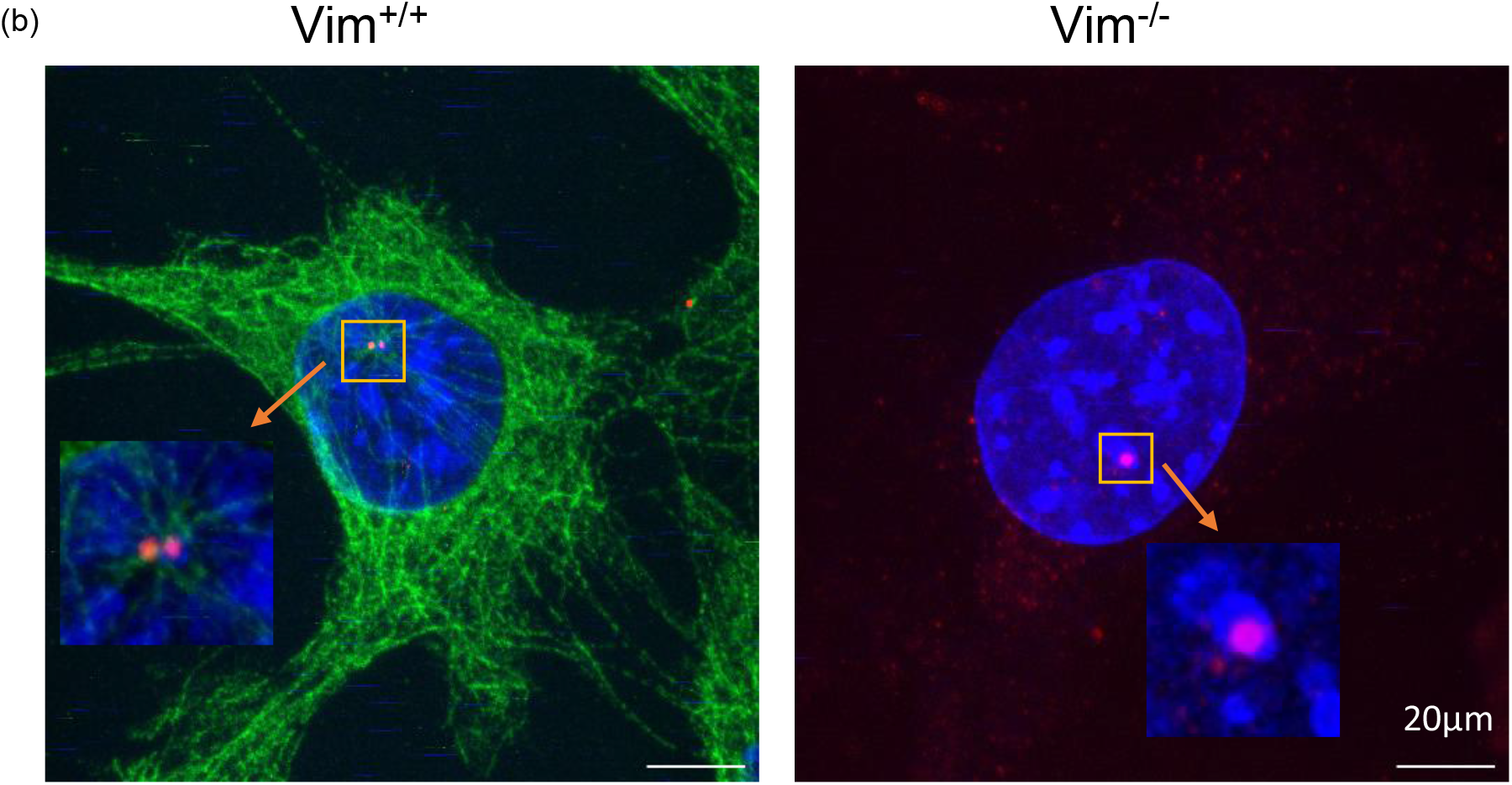
Expansion microscopy images for vim^+/+^ and vim^-/-^ cells stained for nucleus (blue), vimentin (green) and centrosome protein-centrin (red). The inset shows zoomed in images at the centrosome.

### Loss of vimentin disrupts centrosome-mediated microtubule re-nucleation

Cep215, γ-tubulin and pericentrin are PCM proteins associated with the organization and nucleation of microtubules from the centrosome (*30-33*). Thus, based on our results from Fig. 1, we hypothesized that loss of vimentin may disrupt the microtubule nucleation function of the centrosome. To quantitatively analyze centrosome-mediated microtubule-nucleating activity, we performed nocodazole-washout microtubule re-nucleation assays in vim^+/+^ and vim^-/-^ mEFs (Methods, Fig. 3a). Microtubules were first disassembled by treating cells with 1 μM nocodazole for 30 minutes, and then microtubule nucleation from the centrosome was examined at different times after nocodazole washout (0,1, 2 and 5 minutes).

**Figure 3:**
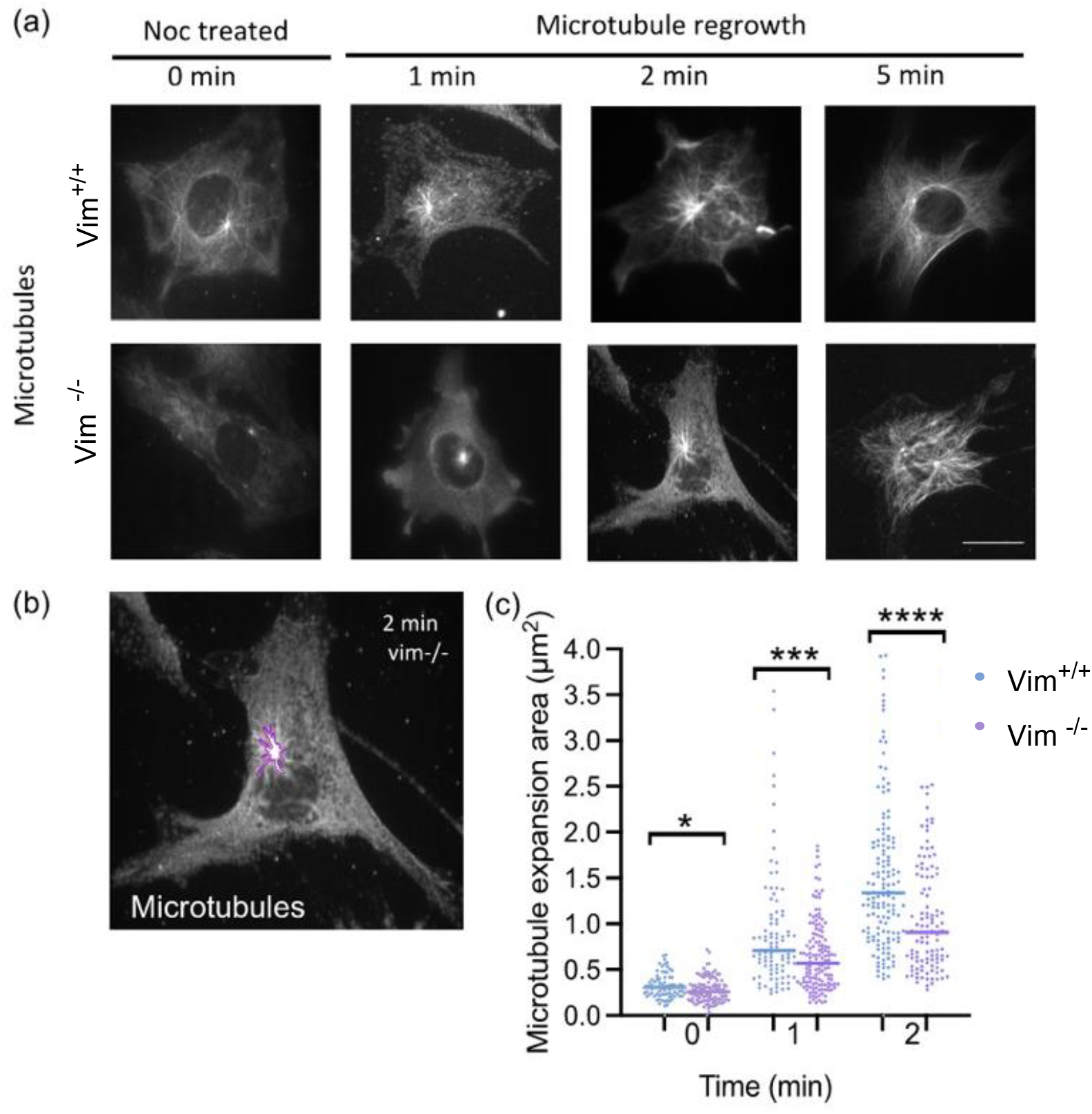
**(a)** Epi-fluorescence microscopy images of microtubules in vim^+/+^ and vim^-/-^ mEFs after nocodazole treatment (0 min) and regrowth at 1, 2, and 5 minutes after nocodazole washout. **(b)** The area marked to study microtubule re-nucleation. **(c)** The quantification of microtubule expansion area in vim^+/+^ and vim^-/-^ mEFs for 0,1 and 2 mins respectively. (Denotation: *, *p* ≤ 0.05; ***, *p* ≤ 0.001; n.s., *p* > 0.05). N ≥ 75 cells analyzed per condition, n=3. The scale bar is 20 μm.

The rate of microtubule re-nucleation was quantified by tracing the area around the centrosome showing microtubules re-nucleation (Fig. 3b). At time = 0 minutes (post nocodazole washout), the amount of tubulin at the centrosome is slightly lower in vimentin-null cells compared to wild-type cells (12% difference, p=0.04). At 1 min of regrowth, the centrosome in vim ^+/+^ showed 28% more microtubule re-nucleation than in null-cells (p<0.001), and by 2 min, regrowth was 33% greater in wild-type than null-cells (p<0.001). At 5 min, re-nucleated microtubules span nearly the whole length of cell for both vim^+/+^ and vim^-/-^ mEFs (Fig 3a). By computing the change in centrosome-nucleated microtubule area over 0-2 minutes, we find that the area of centrosome-nucleated microtubules grows at a rate 40% higher in vim^+/+^ mEFs (0.64 μm^2^/min) than vim^-/-^ mEFs (0.39 μm^2^/min) (Fig. 3c). Taken together, the results in Fig. 3 indicate vimentin impacts centrosome function, in particular enhancing the microtubule-nucleation activity of centrosomes.

### Vimentin increases the amount of acetylated microtubules

On analyzing the nocodazole-treated cells (Fig. 3), we observed the presence of stable microtubule filaments that remained in vim^+/+^ mEFs but not vim ^-/-^ mEFs after sustained nocodazole treatment. As shown in Fig. 4a., after nocodazole treatment (t=0 min), a number of long microtubule filaments persist in wild-type mEF that are not observed in the null cells. To quantify the amount of remaining microtubules, microtubules were identified using a line/curve detection algorithm (Source Steger’s algorithm, Methods) and the total microtubule contour length was computed per cell. As shown in Fig. 4b, wild-type cells had a greater amount of nocodazole-resistant microtubule filaments than vimentin-null cells.

**Figure 4:**
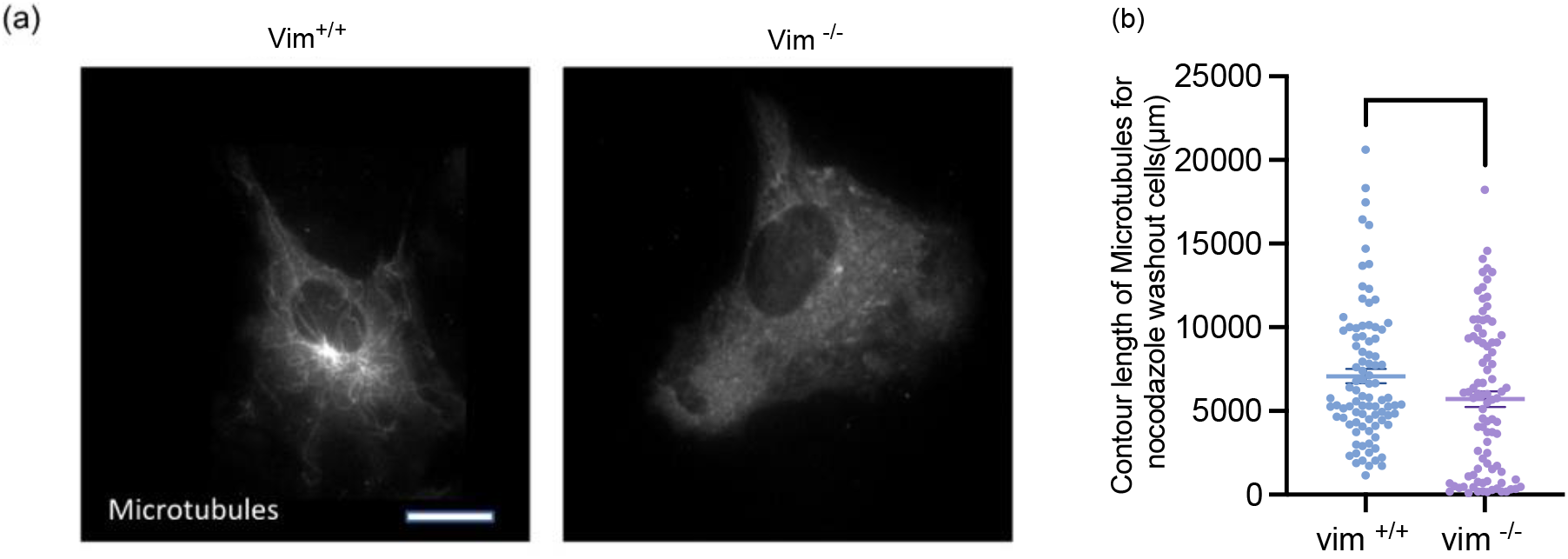
**(a)** Immunofluorescence images of microtubules in nocodazole treated vim ^+/+^ and vim ^-/-^ mEFs. Vim^+/+^ mEF’s show the presence of microtubules even af nocodazole treatment. **(b)** Quantification of the contour length of the microtubules using the curve fit algorithm from FIJI-ImageJ (Denotation: *, *p* ≤ 0.05). N>90 cells analyzed per condition for n=3. The scale bar is 20 μm.

The presence of long-lived nocodazole-resistant microtubules in wild-type mEF indicates a change in microtubule dynamics and stability. Microtubules dynamically assemble and disassemble with a turnover rate of 3-5 minutes (*34*). Nocodazole disrupts microtubule networks by inhibiting microtubule polymerization; for nocodazole-treatment times greater than the microtubule turnover time, microtubules will depolymerize unless they have been stabilized, for instance by post-translational modifications. Acetylation is a post translational modification in which an acetyl group is attached to the lysine group (K40) of α-tubulin (*35*). Acetylation increases the flexibility of microtubules, making them more resistant against mechanical forces and breakage (*35*). In living cells, microtubules are frequently damaged and subsequently depolymerized; thus, acetylation protects against breaking and increases the stability of microtubules (*36*).

To investigate changes in microtubule acetylation levels, we next performed immunofluorescence and immunoblotting studies of acetylated tubulin, as shown in Fig. 5. Figure 5a shows immunofluorescence images of acetylated tubulin in vim ^+/+^ and vim ^-/-^ mEF. In wild-type cells, we observe significant amounts of acetylated tubulin that forms long filaments and a network-like structure in the cell. In contrast, in vimentin-null cells, the acetylated tubulin is much sparser and forms mostly small filamentous squiggles. As shown in Fig. 5b, the mean immunofluorescence intensity of acetylated tubulin is approximately 20% higher in vim^+/+^ cells compared to vim^-/-^ cells (*p* ≤ 0.001;). The immunoblotting studies further confirmed that acetylated tubulin is higher in vim^+/+^ cells compared to vim^-/-^ cells whereas the total alpha-tubulin levels in both the cell lines is the same (Fig 5c and d). Taken together, the results in Fig. 4 and 5 suggest vimentin positively influences the levels of microtubule acetylation, which enhances microtubule stability.

**Figure 5:**
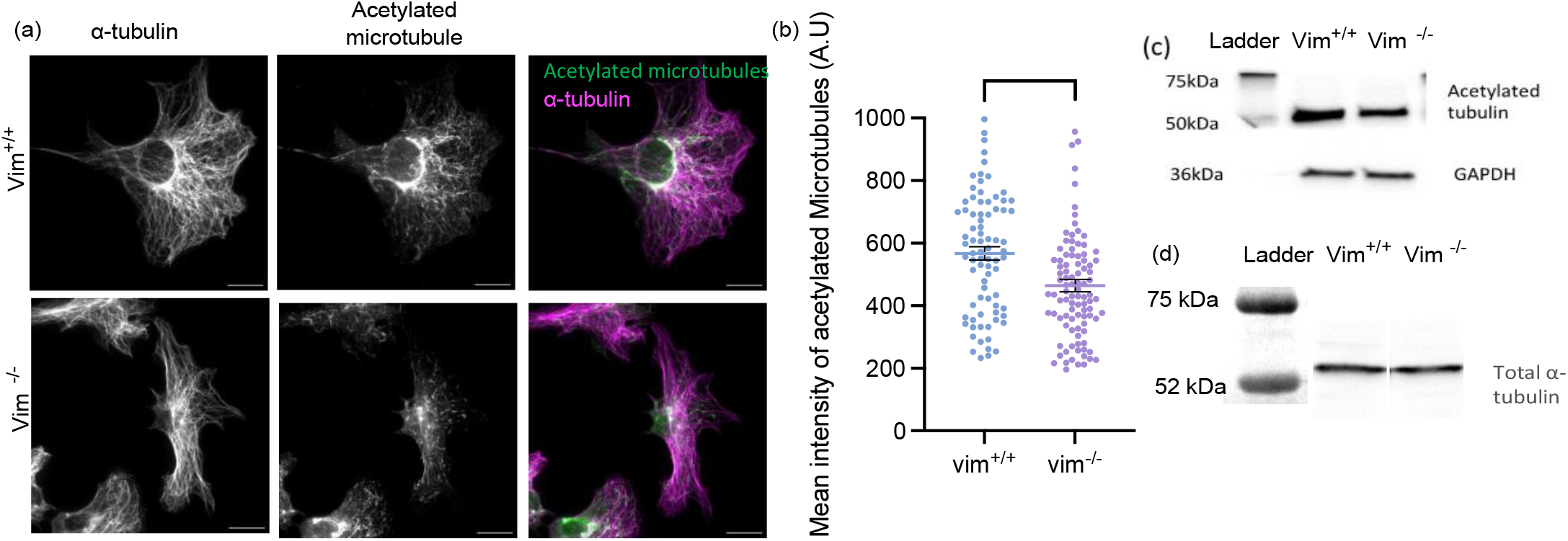
**(a)** Immunofluorescence images of acetylated microtubules for vim^+/+^ and vim^-/-^ mEFs. **(b)** quantification of mean intensity of acetylated microtubules for vim^+/+^ and vim^-/-^ mEFs (Denotation: ***, p ≤ 0.001). N ≥ 90 cells analyzed per condition, n=3. **(c)** Western blot analysis for acetylated tubulin in vim^+/+^ and vim^-/-^ mEFs with GAPDH as control. **(d)** Western blot analysis for total alpha-tubulin in vim+/+ and vim-/- cells. The scale bar is 20 μm.

### Vimentin enhances repositioning of the centrosome towards the wound edge

Cells lacking vimentin have been shown to have decreased cell motility (*22, 37*) As the positioning of the centrosome is important for directional cell polarization and migration, we hypothesized vimentin would also impact the positioning of the centrosome in migrating cells. To study this, we performed an *in-vitro* wound healing assay in vim ^+/+^ and vim ^-/-^ cells. The cells were fixed at 1, 2 and 4 hours after making the scratch and stained for centrosomal protein Cep215, α-tubulin, and the cell nucleus (DAPI) (Fig. 6a).

**Figure 6:**
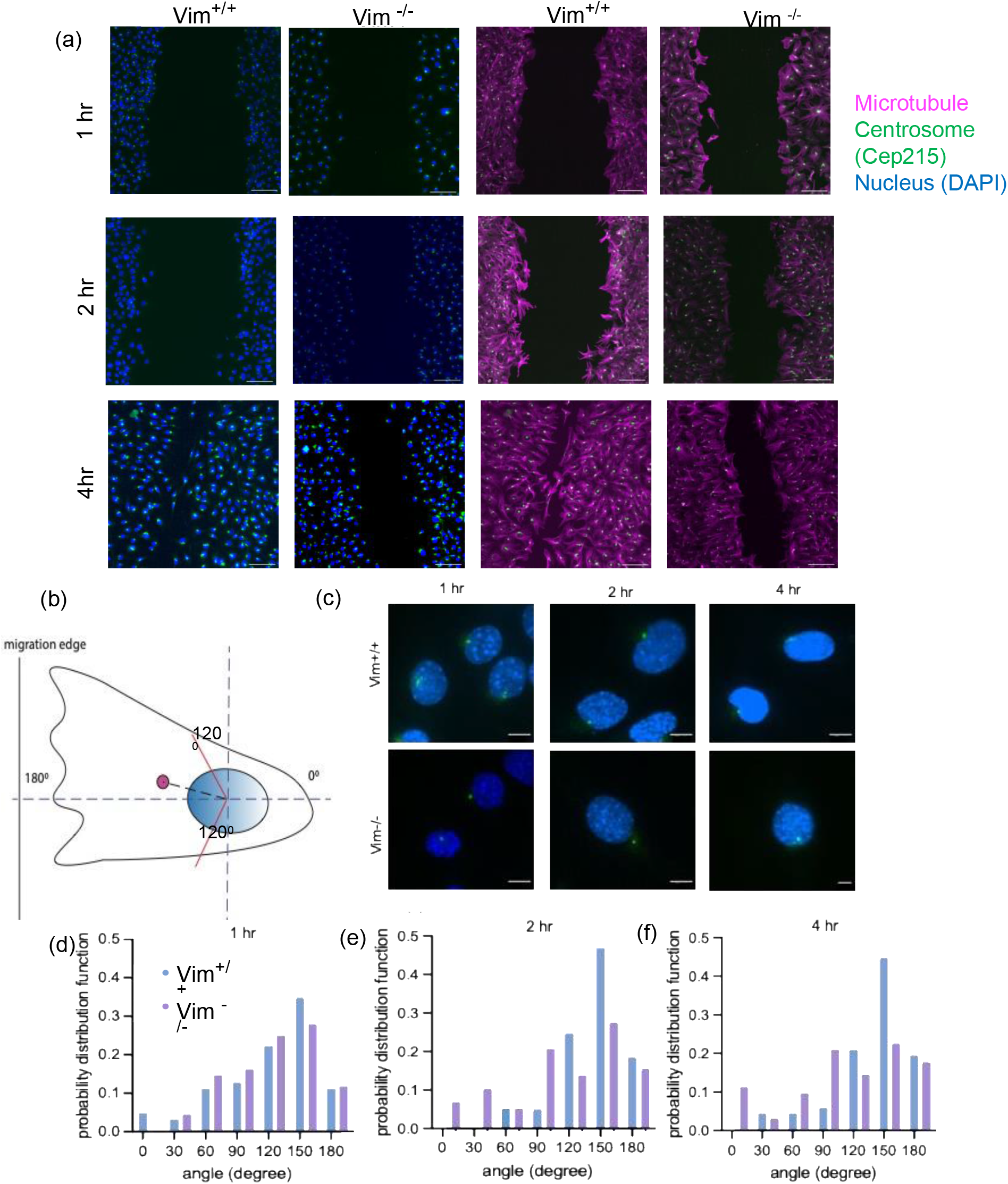
**(a)** Spinning disk confocal images of nucleus (blue), centrosome protein Cep215 (green) and alpha-tubulin (magenta) for vim^+/+^ and vim^-/-^ mEFs fixed at 1,2, and 4 hours respectively after scratch using 10X objective. Scale bar is 100μm. **(b)** Schematic depicting the angle made by the centrosome with respect to the center of the nucleus as the origin axis, the red lines indicate 120^0^ angle **(c)** Zoomed in images (100X objective) of centrosome (green) and nucleus (blue) of vim^+/+^ and vim^-/-^ cells near the wound edge (20 μm from sctrach) at 1,2 and 4 hrs respectively (with scratch to the left of the cell). The scale bar is 10μm **(d-f)** quantification of the centrosome angle for vim^+/+^ and vim^-/-^ cells with respect to the nucleus at 1, 2 and 4 hrs respectively. N ≥ 70 cells analyzed per condition, n=3.

On 2D surfaces, the cell centrosome is localized near the nucleus and its positioning between the cell nucleus and the leading edge of the cell tends to demark the polarized directionality of the cell. Upon wounding, fibroblasts at the edge of the wound become characteristically elongated and polarized with their long axis pointing in the direction of the open wound to facilitate wound healing. To investigate the role of vimentin in cell polarization upon wounding, we quantified the ability of the vim ^+/+^ and vim ^-/-^ mEF to reposition the cell centrosome toward the wound edge. Here, we estimated the cell polarization direction as the vector connecting the center of the cell nucleus to the centrosome and measured its angular position with respect to the direction of the wound edge, with 180° indicating the centrosome was towards the wound and 0° away from the wound edge (Fig. 6b).

Figure 6d-6f shows the probability distribution function (pdf) of centrosome angle for wild-type and vimentin-null fibroblasts near the wound edge (within 20μm from scratch) at 1, 2, and 4 hrs post wounding (Methods). As shown in Fig. 6d, at 1 hr past wounding, both cell types exhibit a polarization toward the wound edge with a peak at approximately 150°, though there is no statistical difference between the two cell types. No significant differences in the centrosome positioning were observed for regions far away from the wound (SI Fig 2). At 2 hr past wounding, the peak at 150° increases from a pdf value of 0.349 to 0.469 for wild-type cells but remains approximately the same for vimentin-null cells. At 4 hours, a peak at 150° remains for wild-type cells, whereas the pdf is leveling out in vimentin-null cells. To further quantify the centrosome positioning, we calculated the percentage of cells with centrosome angles above and below a 120° threshold, with centrosome angle above 120° indicating a polarized direction toward the wound edge (Table 1a). At hr 2 and 4 post wounding, there is a statistically significant increase in polarized cells in wild-type mEF compared to vimentin-null mEF. At 2 hr, approximately 65% of the vim^+/+^ cells are polarized, whereas for the vim^-/-^ cells only 40% of cells showed centrosomes aligned towards the wound (p=0.0018,**). At 4hr, again we see similar results, with 64% vim^+/+^ cells having their centrosomes positioned towards the wound edge, compared to only 40% vim^-/-^ cells (p=0.0007,***). Taken together, the data in Fig. 6 indicate that vimentin enhances cell polarization during wounding healing and supports the repositioning of the centrosome towards the wound edge for directional cell migration to close the wound.

**Table 1:** Percentage of vim^+/+^ and vim^-/-^ cells having centrosome angle above 120^0^ and P-values using Chi-square test for 1, 2 and 4 hours respectively.

| Time (hr) | $\text{Vim}^{+/+}$ | $\text{Vim}^{-/-}$ | P value - Chi-square test |
| --- | --- | --- | --- |
| 1 | 54 | 60 | 0.3915, n.s |
| 2 | 65 | 43 | 0.0018, ** |
| 4 | 64 | 40 | 0.0007, *** |

## Discussion

Coordinated polarized cell migration requires an organized effort of three cytoskeletal networks-F-actin, microtubules, and vimentin but the details of how these distinct networks interact to enable cell motility remain largely unclear. Our results here suggest that vimentin networks are important for centrosome function and stabilizes microtubule dynamics. In the presence of vimentin, we found that the size of centrosome is amplified, and centrosome-mediated microtubule nucleation is enhanced. The presence of vimentin also increases levels of microtubule acetylation, boosting microtubule stability in the cell. Further, during wound healing experiments, vimentin increases repositioning of the cell centrosome toward the wound edge, supporting polarized cell migration to close the wound. From these observations, we propose that vimentin can modulate centrosome structure and function as well as microtubule network stability to enhance cell polarization.

Recent work is revealing intricate, complex pathways at the signaling crossroads among vimentin, microtubules, and F-actin in coordinating cell dynamics. Prior work by Jiu et al indicated that vimentin is antagonistic to actin stress fiber assembly through the microtubule-associated guanine nucleotide exchange factor GEF-H1 that activates RhoA (*38*). FRAP studies showed loss of vimentin increased GEF-H1 dynamics and increased GEF-H1 phosphorylation on Ser886, which was attributed to increased GEF-H1 activity in the absence of vimentin by using a phosphomimetic mutant protein (*38*). Our results here suggest a possible parallel pathway for increased GEF-H1 activity in vimentin-deficient cells through altered microtubule acetylation (Fig 5): we speculate that the increased levels of actin stress fiber assembly and cellular traction stresses in vimentin-deficient cells, which have been observed before, arises from the loss of stable acetylated microtubules, which releases GEF-H1 and increases the soluble pool of GEF-H1 that activates RhoA. On the other hand, recent work on microtubule acetylation by Seetharaman *et al* would predict the reverse effect. Seetharaman *et al* found that microtubule acetylation can promote the release of GEF-H1 from microtubules to activate RhoA and actomyosin contractility (*39*). This indicates a mechanism by which GEF-H1 activity increases with the level of acetylated microtubules, which here is greater in wild-type than vimentin-null mEF and would thus predict an increase in actin stress fiber assembly and contractility in wild-type cells, which we note has been observed in some contexts, such as cells plated on soft substrates or in soft gels compared to on rigid substrates (*37, 40, 41*). These two different expected effects of increased microtubule acetylation in vimentin-expressing cells on RhoA activity through GEF-H1 suggest at least two distinct biochemical signals that engage vimentin in regulating GEF-H1 activity and actin stress fiber formation. Of note, a recent study showed a link between centrosome amplification, an increased number of centrosomes commonly found in cancer cells, with increased tubulin acetylation levels (*42*). Further, the polarized distribution of acetylated tubulin played a role in positioning the cell centrosome, distancing it away from the cell nucleus, and organizing cell polarization.

Our results showing vimentin intermediate filaments impact centrosome size and function might be surprising, given that the primary function of the cell centrosome in organizing microtubules.

Given the interdependent organization of vimentin and microtubules, early work in the intermediate filament field did examine a possible link between vimentin and the centrosome (*43, 44*). Results have been mixed. One of the first studies using electron microscopy in 1985 found no direct interaction between centrioles and vimentin in HeLa cells (*43*), whereas later in 1995 an association of vimentin and pericentriolar material was found in SW-13 cells stably transfected with low levels of vimentin (*44*). Our results indicate a possible functional role of vimentin in the microtubule-nucleating capacities of the centrosome. Our results suggest that vimentin impacts the size of the pericentral material of the cell centrosome, though it is not yet clear whether the change in size is due to altered spatial distribution of centrosomal protein or a change in the amount of PCM protein per centrosome, which are both possible structural changes for the centrosome (*45*). While there are relatively few studies on the association of vimentin and the centrosome, we note that vimentin was identified in a 2003 proteomic characterization of the human centrosome by protein correlation profiling (*46*). Taken together, these works highlight a need to re-examine the link between vimentin, centrosome, and its implications for whole cell organization and polarity.

Our results here suggest new mechanisms by which vimentin coordinates polarized cell migration by regulating centrosome function and microtubule stability. Our recent work has shown loss of vimentin leads to abnormally persistent cell migration through confining microfluidic channels and that the presence of vimentin is needed for the ability of cells to stop and turn around (*23, 24*). In confining three-dimensional spaces, such as tissue, tight spaces add extra constraints and restriction to motion of the cytoskeleton. In many (though not all) 3D settings, the centrosome trails the cell nucleus, defining the tailing direction of the cell, and for the cell to change direction, the centrosome must reposition around the nucleus to a new trailing side of the cell (*47*). We propose that by promoting centrosome activity vimentin boosts the ability of the centrosome to dynamically organize and re-organize microtubules and thus the capacity of the cell to coordinate polarized, directional motion. The microtubule network stabilizing effect of vimentin of the microtubule network may further allow the cell to build an internal compass that competes against externally imposed constraints.

## Conclusions

Taken together, our results demonstrate a new role for VIFs in maintaining centrosome size and microtubule network stability. Our data provides evidence that VIFs are involved in regulating the microtubule-nucleating function of the cell centrosome and the expression levels of acetylated tubulin. Further, during wound healing experiments, vimentin increases repositioning of the cell centrosome toward the wound edge, supporting polarized cell migration to close open wounds. Our results provide new insight into the coordinated crosstalk between vimentin and the microtubule networks in cell polarization.

## Acknowledgements

We thank J.M. Schwarz, Sarthak Gupta, Vladimir Gelfand, Paul Janmey and Annica Gad for useful and insightful discussions. We also thank Jennifer Ross for access and assistance with confocal imaging. This work was supported by and NIH R35 GM142963 awarded to A.E.P and an SU BioInspired Seed Grant awarded to H.H. and A.E.P, and by the National Institutes of Health grants R01GM127621 and R01GM130874 to H.H. Centrosome schematic was created with BioRender.com.

## References

1. D. A. Fletcher, R. D. Mullins, Cell mechanics and the cytoskeleton. Nature 463, 485–492 (2010).

2. M. L. Gardel, I. C. Schneider, Y. Aratyn-Schaus, C. M. Waterman, Mechanical integration of actin and adhesion dynamics in cell migration. Annu Rev Cell Dev Biol 26, 315–333 (2010).

3. M. Murrell, P. W. Oakes, M. Lenz, M. L. Gardel, Forcing cells into shape: the mechanics of actomyosin contractility. Nature Reviews Molecular Cell Biology 16, 486–498 (2015).

4. M. Kikumoto, M. Kurachi, V. Tosa, H. Tashiro, Flexural Rigidity of Individual Microtubules Measured by a Buckling Force with Optical Traps. Biophysical Journal 90, 1687–1696 (2006).

5. J. C. M. Meiring, B. I. Shneyer, A. Akhmanova, Generation and regulation of microtubule network asymmetry to drive cell polarity. Current Opinion in Cell Biology 62, 86–95 (2020).

6. S. Stehbens, T. Wittmann, Targeting and transport: How microtubules control focal adhesion dynamics. Journal of Cell Biology 198, 481–489 (2012).

7. A. E. Patteson et al., Vimentin protects cells against nuclear rupture and DNA damage during migration. J Cell Biol 218, 4079–4092 (2019).

8. M. Guo et al., The Role of Vimentin Intermediate Filaments in Cortical and Cytoplasmic Mechanics. Biophysj 105, 1562–1568 (2013).

9. Z. Gan et al., Vimentin Intermediate Filaments Template Microtubule Networks to Enhance Persistence in Cell Polarity and Directed Migration. Cell Syst 3, 252-263.e258 (2016).

10. I. Dupin, Y. Sakamoto, S. Etienne-Manneville, Cytoplasmic intermediate filaments mediate actin-driven positioning of the nucleus. Journal of Cell Science 124, 865–872 (2011).

11. F. Huber, A. Boire, M. P. López, G. H. Koenderink, Cytoskeletal crosstalk: when three different personalities team up. Current opinion in cell biology 32, 39–47 (2015).

12. S. Seetharaman, S. Etienne-Manneville, Cytoskeletal Crosstalk in Cell Migration. Trends Cell Biol 30, 720–735 (2020).

13. M. Yoon, R. D. Moir, V. Prahlad, R. D. Goldman, Motile properties of vimentin intermediate filament networks in living cells. The Journal of cell biology 143, 147–157 (1998).

14. L. Schaedel, C. Lorenz, A. V. Schepers, S. Klumpp, S. Köster, Vimentin intermediate filaments stabilize dynamic microtubules by direct interactions. Nature Communications 12, 3799 (2021).

15. R. D. Goldman, S. Khuon, Y. H. Chou, P. Opal, P. M. Steinert, The function of intermediate filaments in cell shape and cytoskeletal integrity. J Cell Biol 134, 971–983 (1996).

16. M. G. Mendez, S.-I. Kojima, R. D. Goldman, Vimentin induces changes in cell shape, motility, and adhesion during the epithelial to mesenchymal transition. FASEB journal : official publication of the Federation of American Societies for Experimental Biology 24, 1838–1851 (2010).

17. A. E. Patteson, A. Vahabikashi, R. D. Goldman, P. A. Janmey, Mechanical and NonMechanical Functions of Filamentous and Non-Filamentous Vimentin. Bioessays 11, e2000078 (2020).

18. A. Satelli, S. Li, Vimentin in cancer and its potential as a molecular target for cancer therapy. Cell Mol Life Sci 68, 3033–3046 (2011).

19. A. Johnson, J. Lewis, B. Alberts, Molecular biology of the cell. (2002).

20. G. W. Luxton, G. G. Gundersen, Orientation and function of the nuclear-centrosomal axis during cell migration. Curr Opin Cell Biol 23, 579–588 (2011).

21. S. H. Shabbir, M. M. Cleland, R. D. Goldman, M. Mrksich, Geometric control of vimentin intermediate filaments. Biomaterials 35, 1359–1366 (2014).

22. B. T. Helfand et al., Vimentin organization modulates the formation of lamellipodia. Molecular biology of the cell 22, 1274–1289 (2011).

23. A. E. Patteson et al., Loss of Vimentin Enhances Cell Motility through Small Confining Spaces. Small 15, e1903180 (2019).

24. S. Gupta, A. E. Patteson, J. M. Schwarz, The role of vimentin–nuclear interactions in persistent cell motility through confined spaces. New journal of physics 23, 093042 (2021).

25. N. Costigliola et al., Vimentin fibers orient traction stress. Proceedings of the National Academy of Sciences 114, 5195–5200 (2017).

26. M. T. H. Thanh, A. Grella, D. Kole, S. Ambady, Q. Wen, Vimentin intermediate filaments modulate cell traction force but not cell sensitivity to substrate stiffness. Cytoskeleton, (2021).

27. N. Krishnan et al., Rab11 endosomes and Pericentrin coordinate centrosome movement during pre-abscission in vivo. Life science alliance 5, (2022).

28. A. Aljiboury et al., Pericentriolar matrix (PCM) integrity relies on cenexin and polo-like kinase (PLK) 1. Molecular biology of the cell 33, br14 (2022).

29. C. Steger, Unbiased extraction of lines with parabolic and Gaussian profiles. Computer Vision and Image Understanding 117, 97–112 (2013).

30. S. Graser, Y. D. Stierhof, E. A. Nigg, Cep68 and Cep215 (Cdk5rap2) are required for centrosome cohesion. Journal of Cell Science 120, 4321–4331 (2007).

31. K.-W. Fong, Y.-K. Choi, J. B. Rattner, R. Z. Qi, CDK5RAP2 Is a Pericentriolar Protein That Functions in Centrosomal Attachment of the γ-Tubulin Ring Complex. Molecular Biology of the Cell 19, 115–125 (2008).

32. J. B. Dictenberg et al., Pericentrin and γ-Tubulin Form a Protein Complex and Are Organized into a Novel Lattice at the Centrosome. Journal of Cell Biology 141, 163–174 (1998).

33. T. Stearns, L. Evans, M. Kirschner, GAMMA-TUBULIN IS A HIGHLY CONSERVED COMPONENT OF THE CENTROSOME. Cell 65, 825–836 (1991).

34. E. Shelden, P. Wadsworth, Observation and quantification of individual microtubule behavior in vivo: microtubule dynamics are cell-type specific. The Journal of cell biology 120, 935–945 (1993).

35. C. Janke, G. Montagnac, Causes and consequences of microtubule acetylation. Current Biology 27, R1287–R1292 (2017).

36. Z. Xu et al., Microtubules acquire resistance from mechanical breakage through intralumenal acetylation. Science 356, 328–332 (2017).

37. B. Eckes et al., Impaired mechanical stability, migration and contractile capacity in vimentin-deficient fibroblasts. Journal of cell science 111, 1897–1907 (1998).

38. Y. Jiu et al., Vimentin intermediate filaments control actin stress fiber assembly through GEF-H1 and RhoA. J Cell Sci 130, 892–902 (2017).

39. S. Seetharaman et al., Microtubules tune mechanosensitive cell responses. Nature Materials 21, 366–377 (2022).

40. F. Alisafaei et al., Vimentin Intermediate Filaments Can Enhance or Abate Active Cellular Forces in a Microenvironmental Stiffness-Dependent Manner. bioRxiv, 2022.2004.2002.486829 (2022).

41. A. Vahabikashi et al., Probe Sensitivity to Cortical versus Intracellular Cytoskeletal Network Stiffness. Biophys J 116, 518–529 (2019).

42. P. Monteiro, B. Yeon, S. S. Wallis, S. A. Godinho, Centrosome amplification fine tunes tubulin acetylation to differentially control intracellular organization. The EMBO Journal 42, e112812 (2023).

43. B. Maro, M. Paintrand, M. E. Sauron, D. Paulin, M. Bornens, Vimentin filaments and centrosomes. Are they associated? Exp Cell Res 150, 452–458 (1984).

44. K. T. Trevor, J. G. McGuire, E. V. Leonova, Association of vimentin intermediate filaments with the centrosome. J Cell Sci 108 (Pt 1), 343–356 (1995).

45. D. G. Sankaran, A. J. Stemm-Wolf, B. L. McCurdy, B. Hariharan, C. G. Pearson, A semiautomated machine learning-aided approach to quantitative analysis of centrosomes and microtubule organization. J Cell Sci 133, (2020).

46. J. S. Andersen et al., Proteomic characterization of the human centrosome by protein correlation profiling. Nature 426, 570–574 (2003).

47. J. Zhang, Y.-l. Wang, Centrosome defines the rear of cells during mesenchymal migration. Molecular biology of the cell 28, 3240–3251 (2017).

