## Supplemental fig1 and fig6 for "Vimentin supports cell polarization by enhancing centrosome function and microtubule acetylation"

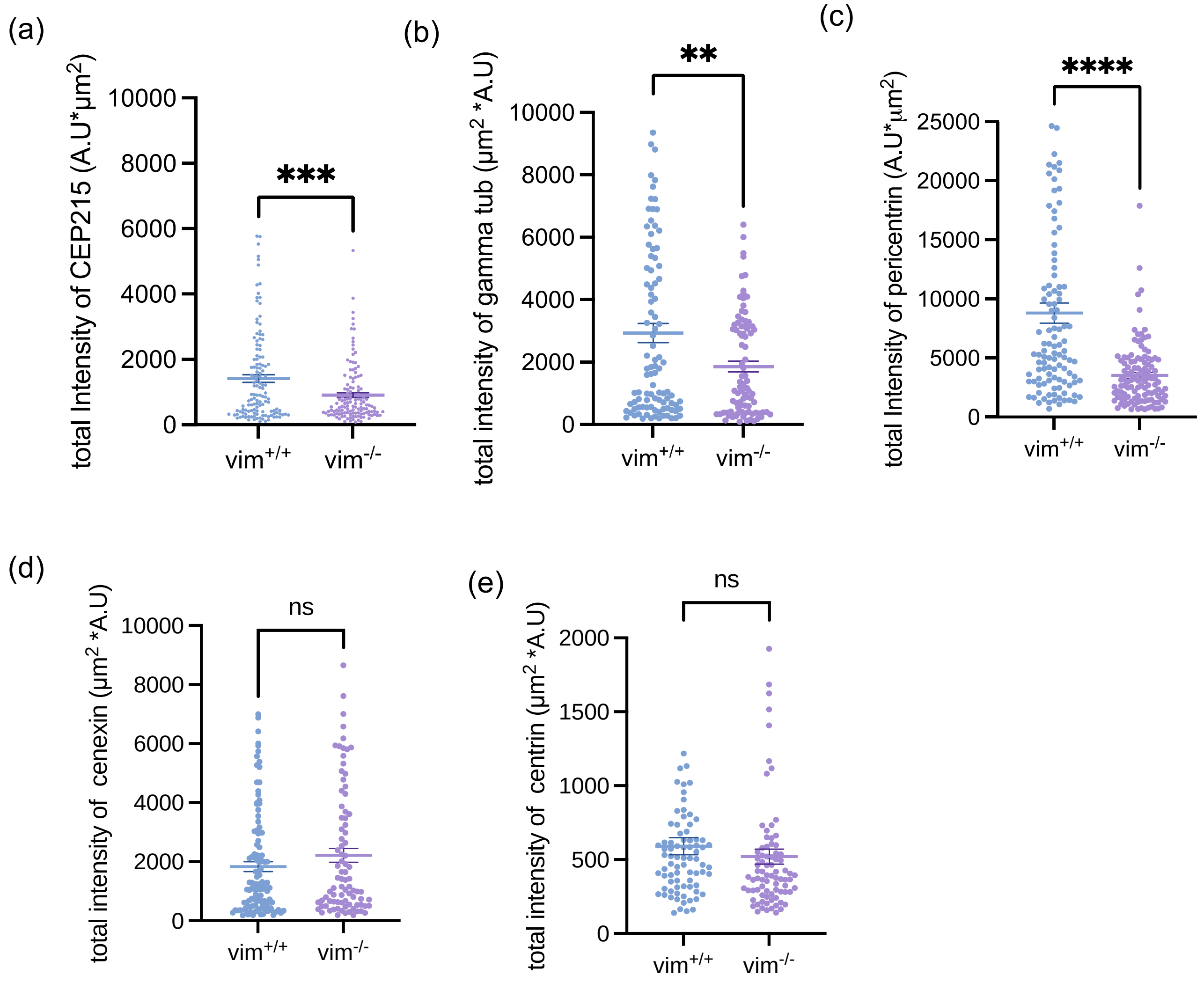


**Figure S1 (a-e):** Total intensity plots for vim$^{+/+}$ and vim$^{-/-}$ cells stained for CEP215, gamma tubulin, pericentrin, cenexin and centrin. Denotation: ***, $p \leq 0.001$; **, $p \leq 0.01$) n=3, N>90 cells analyzed per condition.

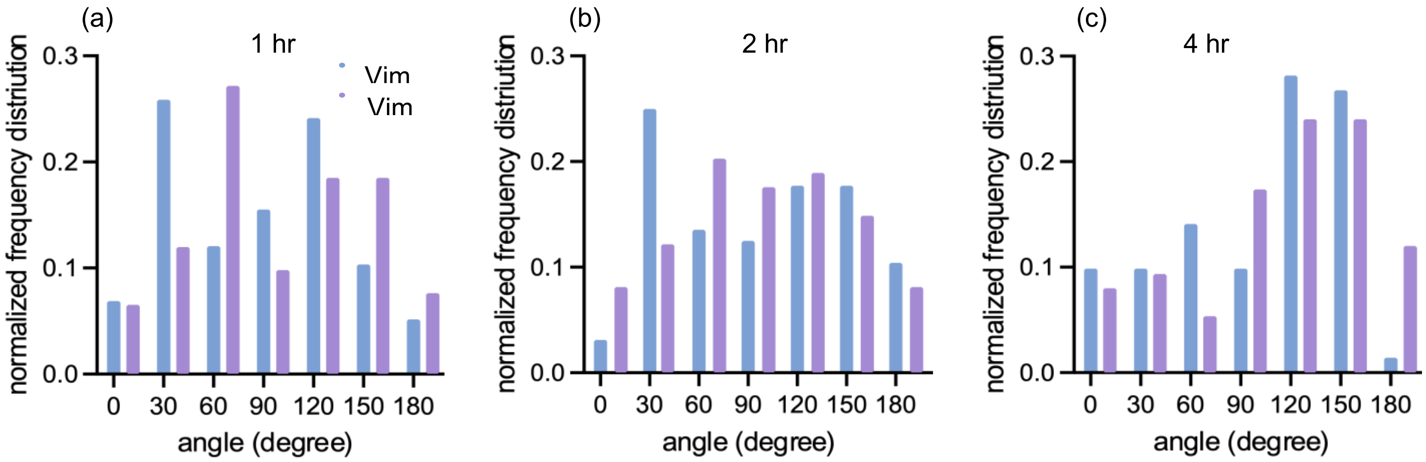

**Figure S2: (a-c)** Quantification of the centrosome angle for $vim^{+/+}$ and $vim^{-/-}$ cells far away from the wound edge (distance greater than 50μm from wound edge) with respect to the nucleus at 1, 2 and 4 hrs respectively.
